# Disabling Cas9 by an anti-CRISPR DNA mimic

**DOI:** 10.1101/129627

**Authors:** Jiyung Shing, Fuguo Jiang, Jun-Jie Liu, Nicholas L. Bray, Benjamin J. Rauch, Seung Hyun Baik, Eva Nogales, Joseph Bondy-Denomy, Jacob E. Corn, Jennifer A. Doudna

**Affiliations:** Innovative Genomics Institute, Berkeley, California 94720; Department of Molecular and Cell Biology, University of California, Berkeley, California 94720; Department of Microbiology and Immunology, and Quantitative Biosciences Institute, University of California, San Francisco, CA 94158; Howard Hughes Medical Institute, Berkeley, California 94720; Department of Chemistry, University of California, Berkeley, California 94720; Molecular Biophysics and Integrated Bioimaging Division, Lawrence Berkeley National Laboratory, Berkeley, California 94720

## Abstract

CRISPR-Cas9 gene editing technology is derived from a microbial adaptive immune system, where bacteriophages are often the intended target. Natural inhibitors of CRISPR-Cas9 enable phages to evade immunity and show promise in controlling Cas9-mediated gene editing in human cells. However, the mechanism of CRISPR-Cas9 inhibition is not known and the potential applications for Cas9 inhibitor proteins in mammalian cells has not fully been established. We show here that the anti-CRISPR protein AcrIIA4 binds only to assembled Cas9-single guide RNA (sgRNA) complexes and not to Cas9 protein alone. A 3.9 Å resolution cryo-EM structure of the Cas9-sgRNA-AcrIIA4 complex revealed that the surface of AcrIIA4 is highly acidic and binds with 1:1 stoichiometry to a region of Cas9 that normally engages the DNA protospacer adjacent motif (PAM). Consistent with this binding mode, order-of-addition experiments showed that AcrIIA4 interferes with DNA recognition but has no effect on pre-formed Cas9-sgRNA-DNA complexes. Timed delivery of AcrIIA4 into human cells as either protein or expression plasmid allows on-target Cas9-mediated gene editing while reducing off-target edits. These results provide a mechanistic understanding of AcrIIA4 function and demonstrate that inhibitors can modulate the extent and outcomes of Cas9-mediated gene editing.

---

Phage-encoded inhibitors of CRISPR-Cas bacterial immune systems evolved to enable phage escape from destruction in bacterial cells (1) and have the potential to control CRISPR-Cas enzymes that are deployed for gene editing applications in various cell types (2, 3). Determination of the molecular basis for Cas9 inhibition could shed light on the evolutionary “arms race” between phage and bacteria, as well as suggest new approaches to regulate genome editing in eukaryotic cells. The 87 amino acid anti-CRISPR-Cas9 protein AcrIIA4 is notable in both respects: it inhibits multiple Cas9 proteins including the widely used Cas9 ortholog from *Streptococcus pyogenes* (Spy), and it blocks Cas9-mediated gene editing in human cells (3). To investigate the molecular basis for AcrIIA4-mediated Cas9 inhibition, we first tested whether recombinant AcrIIA4 protein interacts directly with SpyCas9 (Fig. 1A). Purified AcrIIA4 was incubated with *Spy*Cas9 in the presence or absence of a single-guide RNA (sgRNA) that assembles with Cas9 to provide sequence-specific DNA recognition (4). Size exclusion chromatography showed that AcrIIA4 binds to SpyCas9 only in the presence of sgRNA (Fig. 1B and C, fig. S1), implying that AcrIIA4 recognizes a protein surface created upon sgRNA-triggered conformational rearrangement (5, 6). Furthermore, the AcrIIA4-bound Cas9-sgRNA complex is more resistant to proteolytic digestion than the Cas9-sgRNA complex alone, implying that AcrIIA4 stabilizes a particular Cas9 conformation (fig. S2).

**Fig. 1.**
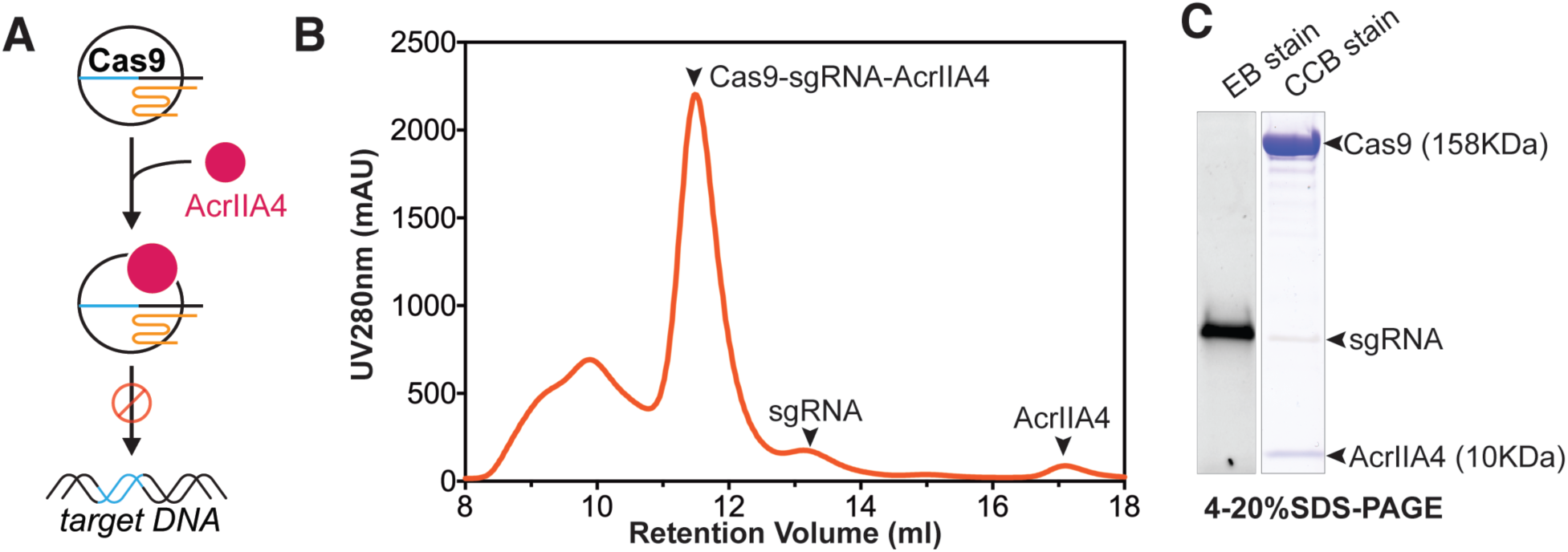
AcrIIA4 binds to the *S. pyogenes* Cas9-sgRNA complex. A) A cartoon depiction of Cas9 protein loaded with the sgRNA binding to AcrIIA4 (pink). Cas9-sgRNA complexed with AcrIIA4 is unable to bind to the target DNA. B) Size exclusion chromatogram of SpyCas9-sgRNA after preincubation with AcrIIA4. Relevant peaks are indicated with arrowheads. C) Coomassie and Ethidium Bromide (EB) stained polyacrylamide gel showing the commigration of AcrIIA4 with Cas9 in the presence of guide RNA.

To elucidate the molecular basis of AcrIIA4-mediated inhibition of Cas9 activity, we performed cryo-electron microscopy (cryo-EM) single-particle analysis on a SpyCas9-sgRNA complex bound to AcrIIA4. Cryo-EM images were collected on a Krios microscope using zero-loss energy-filtered imaging and a K2 direct electron detector. After unsupervised 3D classification of 840,000 particle images, refinement of a class containing 285,600 particles resulted in an EM reconstruction of the SpyCas9-sgRNA-AcrIIA4 complex with an overall resolution of 3.9 Å (fig. S3-S4). Subsequent local 3D classification that was focused on Cas9’s HNH nuclease domain ultimately yielded two EM reconstructions, one at 3.9 Å resolution where the HNH was poorly resolved due to flexibility (reconstruction 1, obtained as a combination of two 3D classes) (Fig. 2a and fig. S5a) and one with better defined density for the HNH domain at an average resolution of 4.5 Å (reconstruction 2) (fig. S4d and S5a; see Methods section and fig. S3). In both cryo-EM reconstructions, the density for the HNH domain of Cas9 was weaker than for the rest of the structure, consistent with the previously observed conformational plasticity of the HNH nuclease domain in the pre-targeting state. An atomic model for AcrIIA4 was built from reconstruction 1. The EM density map displays excellent main-chain connectivity and side-chain densities for almost all residues of AcrIIA4 (fig. S5). We then built an atomic model for the Cas9-sgRNA complex based on the crystal structure of SpyCas9-sgRNA (PDB ID 4ZT0) and refined the entire SpyCas9-sgRNA-AcrIIA4 model in real space to good stereochemistry (Fig. 2B and table S1). Overall, Cas9 bound to AcrIIA4 resembles the pre-target state rather than the DNA-bound state (fig. S6).

**Fig. 2.**
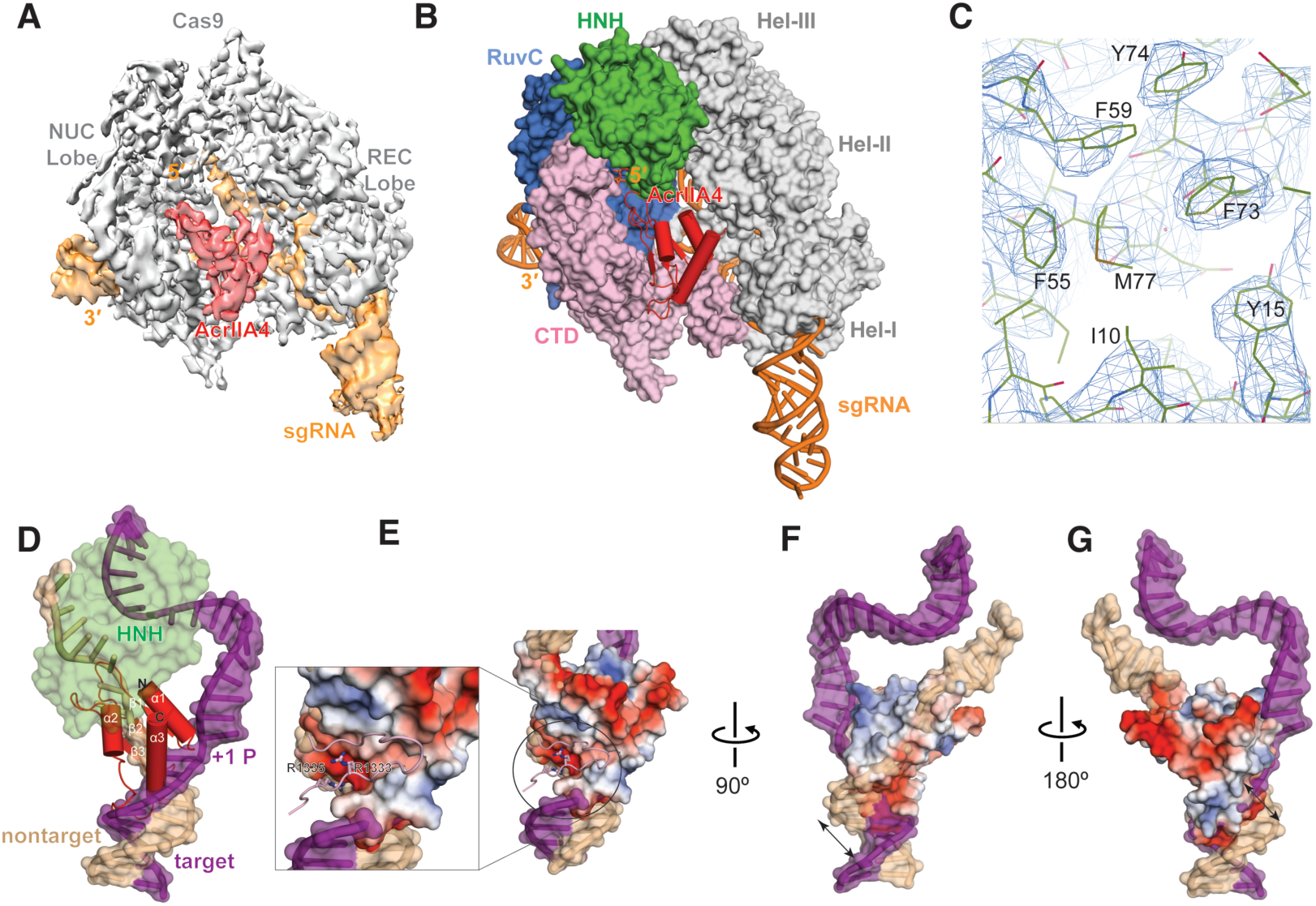
Architecture of the *S. pyogenes* Cas9-sgRNA in complex with AcrIIA4. A) Cryo-EM reconstruction of the AcrII4-bound SpyCas9. The electron density map was contoured at high threshold levels showing distinct features for each subunit. B) The atomic model of SpyCas9-sgRNA-ArcrIIA4. AcrIIA4 (red) and sgRNA (orange) are shown in ribbon diagram and SpyCas9 is displayed as surface representation. C) Representative region (hydrophobic core) of the cryo-EM density for AcrIIA4 with the refined model superimposed. D) Superposition with Cas9-guideRNA-dsDNA structure (PDB ID 5F9R). For clarity, Cas9 is omitted except the HNH domain. Target and nontarget DNA strand is is colored purple and beige, respectively. E) Electrostatic surface potential of AcrIIA4. The inset shows the PAM recognition residues (R1333 and R1334) are largely buried in an acidic pocket within AcrIIA4. F) and G) Close-up of AcrIIA4 binding pocket at different views showing AcrIIA4 is a dsDNA mimic.

*De novo* model building demonstrated that AcrIIA4 binds to Cas9 with 1:1 stoichiometry and comprises a three-stranded antiparallel β sheet flanked by one alpha-helix at the amino-terminal end and two α helices on the carboxy-terminal end (α_1_β_1_β_2_β_3_α_2_α_3_). Remarkably, superposition of the AcrIIA4-bound Cas9 structure with DNA-bound Cas9 revealed that AcrIIA4 sits exactly in the PAM-interacting cleft formed between the alpha-helical recognition (REC) lobe and the nuclease lobe (Fig. 2B and fig. S6). AcrIIA4 completely occupies the PAM binding pocket and thus blocks DNA recognition through contacts between the β3 strand of AcrIIA4 and Cas9 PAM-binding residues (R1333 and R1335) (Fig. 2D-E). In addition, AcrIIA4 wedges into the DNA melting region immediately upstream of the PAM sequence and sits on top of +1 phosphate on the target strand and the flipped nucleotides on the nontarget strand, indicating AcrIIA4 could also prevent DNA binding/unwinding. Consistent with this DNA-mimicing binding mode, AcrIIA4 is extremely acidic (Fig. 2E). Interestingly, AcrIIA4 also occupies the same space as the DNA-bound HNH domain and the linker connecting the HNH and RuvC domains (Fig. 2D), suggesting AcrIIA4 could block the HNH movement required for catalysis (Fig. 2B, C).

Collectively, our structural studies show that AcrIIA4 is a highly acidic DNA mimic that blocks target DNA recognition through multiple mechanisms: 1) competitive inhibition of PAM binding; 2) inhibition of DNA unwinding upstream of the PAM sequence; and 3) inactivation of HNH domain movement from the inactive to active conformation. These structural findings help explain the effectiveness of AcrIIA4 as an inhibitor of Cas9-mediated DNA cleavage and cell-based genome editing. Previous studies revealed the importance of Cas9’s interactions with targeted DNA, mediated via the PAM in an interaction preceding base pairing between the sgRNA and its target (7). Transient Cas9-sgRNA association with PAM sequences in DNA is thought to enable a rapid target sequence search. By mimicking the structure and electrostatic properties of the DNA PAM sequence, AcrIIA4 might compete for initial DNA binding and thereby prevent target recognition and cleavage (Fig. 2E-G).

To determine the functional effects of AcrIIA4 binding to Cas9-sgRNA complexes, we subjected DNA substrates possessing a target sequence to Cas9-sgRNA-catalyzed DNA cleavage assays. A linearized plasmid possessing a target sequence and PAM was incubated with combinations of Cas9, sgRNA and AcrIIA4, and the products were resolved by gel electrophoresis. Cas9-sgRNA alone completely cut the target DNA within five minutes, while AcrIIA4 limited cleavage at even later time points (Fig. 3A). To confirm that AcrIIA4 would also inactivate Cas9 at low enzyme and inhibitor concentrations, we performed similar experiments with a radiolabeled DNA target for increased sensitivity (Fig. 3B). Near-stoichiometric concentrations of AcrIIA4 inhibited DNA cleavage when titrated against 10 nM Cas9-gRNA complex, indicating an apparent dissociation constant of less than 10 nM. These data show that AcrIIA4 functions as a robust Cas9 “off-switch” that can inhibit most Cas9 activity at low concentrations.

**Fig. 3.**
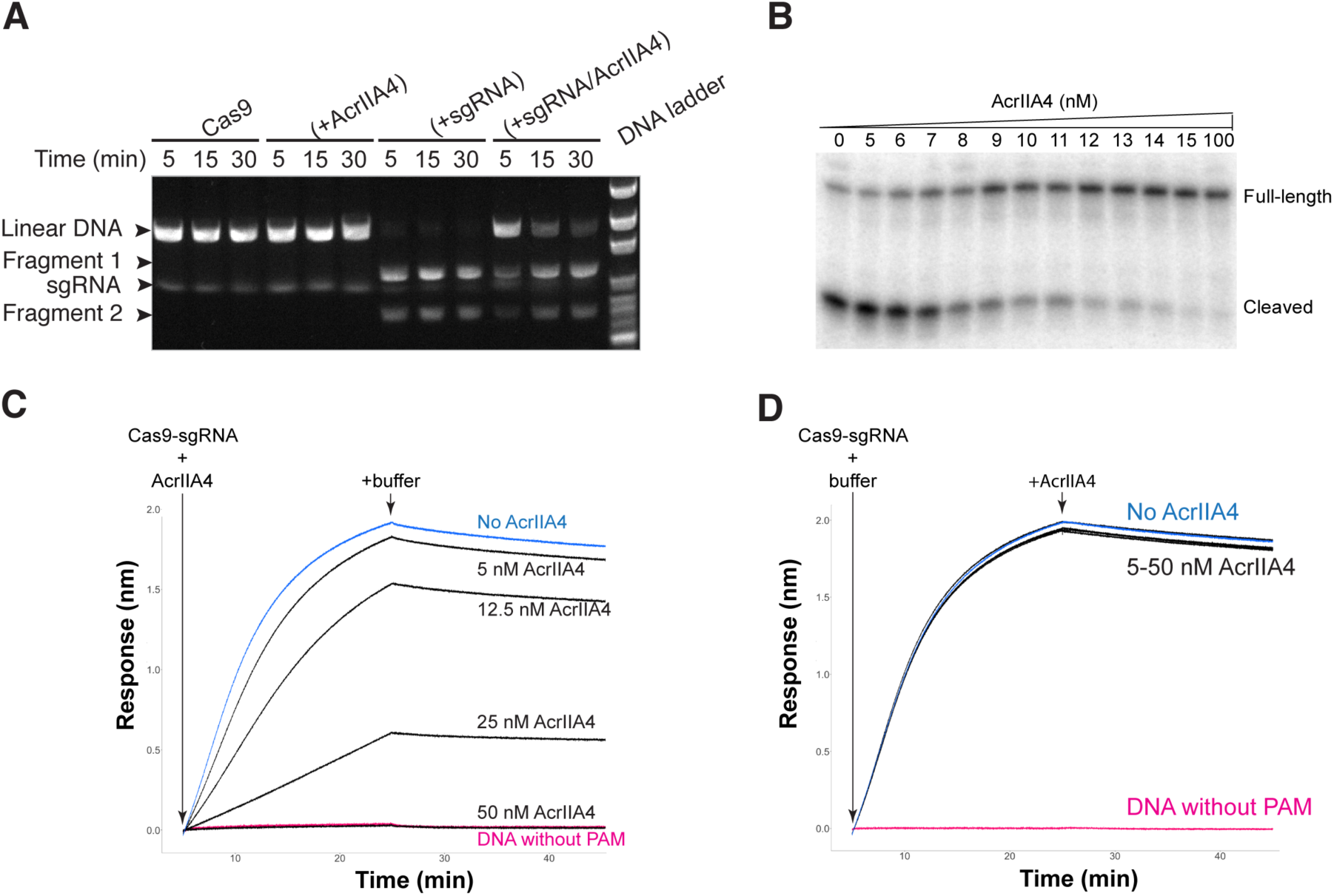
AcrIIA4 inhibits DNA cleavage *in vitro*. A) A linearized plasmid was incubated with Cas9 alone, AcrIIA4 alone, Cas9 + sgRNA, or Cas9 + sgRNA + AcrIIA4 for 5, 15, or 30 minutes. Reactions were resolved via agarose gel electrophoresis and visualized by ethidium bromide staining. B) AcrIIA4 inhibits SpyCas9 cleavage of radiolabeled target DNA *in vitro*. 10 nM SpyCas9-crRNA-tracrRNA complex were pre-incubated with increasing concentrations (0-100 nM) of AcrIIA4. Sub-stoichiometric synthetic oligonucleotide duplexes (2.5 nM) bearing a radiolabel at the 5’ end of the complementary strand, were introduced for 6-minute cleavage reactions. Reactions were resolved by denaturing polyacrylamide gel electrophoresis and visualized by phosphorimaging. C) AcrIIA4 inhibits dCas9-sgRNA binding to a DNA target but does not affect target release, as measured by biolayer interferometry. Pre-incubating increasing concentrations of AcrIIA4 with dCas9-sgRNA reduces the on-rate of association with a DNA target relative to no inhibitor (blue). Maximal inhibition is identical to dCas9-sgRNA added to target DNA with a PAM mutation (red). D) Addition of increasing concentrations of AcrIIA4 with pre-formed dCas9-sgRNA-DNA complex has no effect upon the off-rate of dissociation.

The structure of Cas9 bound to AcrIIA4 suggested that the inhibitor competes for the initial DNA recognition event of PAM binding. Yet we previously found that Cas9 binds so tightly to target DNA that its off-rate is negligible (7, 8). This implies an unusual non-equilibrium mode of inhibition, in which AcrIIA4 requires access to Cas9-sgRNA prior to formation of the Cas9-sgRNA-DNA complex. To test this prediction, we used bilayer interferometry to measure the binding of catalytically inactivated Cas9 (dCas9) to a DNA target in the presence of inhibitor. These experiments were performed under stoichiometric binding conditions in which Cas9 was present at concentrations greater than the dissociation constant of the Cas9-AcrIIA4 interaction. Pre-incubation of Cas9 with AcrIIA4 markedly inhibited Cas9 binding to DNA (on-rate) in a dose-dependent fashion, including complete prevention of target engagement (Fig. 3C, fig. S7). An electrophoretic mobility shift assay (EMSA) further confirmed that AcrIIA4 does not impact sgRNA loading into Cas9 protein alone (fig. S8), but does inhibit Cas9-sgRNA complex binding to a target DNA (fig. S9). Strikingly, allowing the Cas9-sgRNA-DNA complex to form and then adding AcrIIA4 had no effect on Cas9’s release of target DNA (off-rate). These order-of-addition results are consistent with both the structure of AcrIIA4 bound to Cas9 and Cas9’s extremely slow dissociation from DNA. Together, these data support a mechanism for AcrIIA4 inhibition in which the inhibitor blocks Cas9’s ability to bind and cut target DNA by obscuring the PAM-interacting domain.

To determine the ability of AcrIIA4 to regulate gene editing in human cells, we first utilized human K562 erythroleukemia cells stably expressing a chromosomally-integrated blue fluorescent protein (BFP) reporter (8). Nucleofection of Cas9-sgRNA (Cas9 RNP) complexes targeting BFP resulted in loss of BFP fluorescence in almost all cells as measured by flow cytometry (fig. S10). Simultaneous delivery of Cas9 RNP and AcrIIA4 protein inhibited Cas9-mediated gene targeting by up to 80% (Fig. 4A blue symbols; fig. S10). Simultaneous delivery of AcrIIA4 encoded in a plasmid inhibited gene editing to a lesser extent, possibly due to a delay in expression of the inhibitor from a plasmid relative to immediate nucleofection of Cas9 RNP (Fig. 4A green symbols; fig. S10, S11). This implied that the order-of-addition effects we observed in biophysical experiments may play a role during gene editing. To test this idea, we nucleofected AcrIIA4 plasmid 24 hours prior to introducing Cas9 RNP into cells. The presence of the AcrIIA4-encoding plasmid 24 hours prior to Cas9 RNP introduction inhibited Cas9-mediated gene targeting to a far greater extent than was observed during co-introduction, and to a similar extent as observed during co-introduction of AcrIIA4 protein and Cas9 RNP (Fig. 4B, fig. S12). Remarkably, we found that addition of AcrIIA4 six hours after Cas9-RNP reduced editing by ∼50%, demonstrating the utility of inhibitors for revealing *in vivo* gene editing kinetics (Fig. 4C, fig. S13). In sum, controlling the timing of AcrIIA4 inhibition in human cells strongly affects the frequency of gene editing at a given locus.

**Fig. 4.**
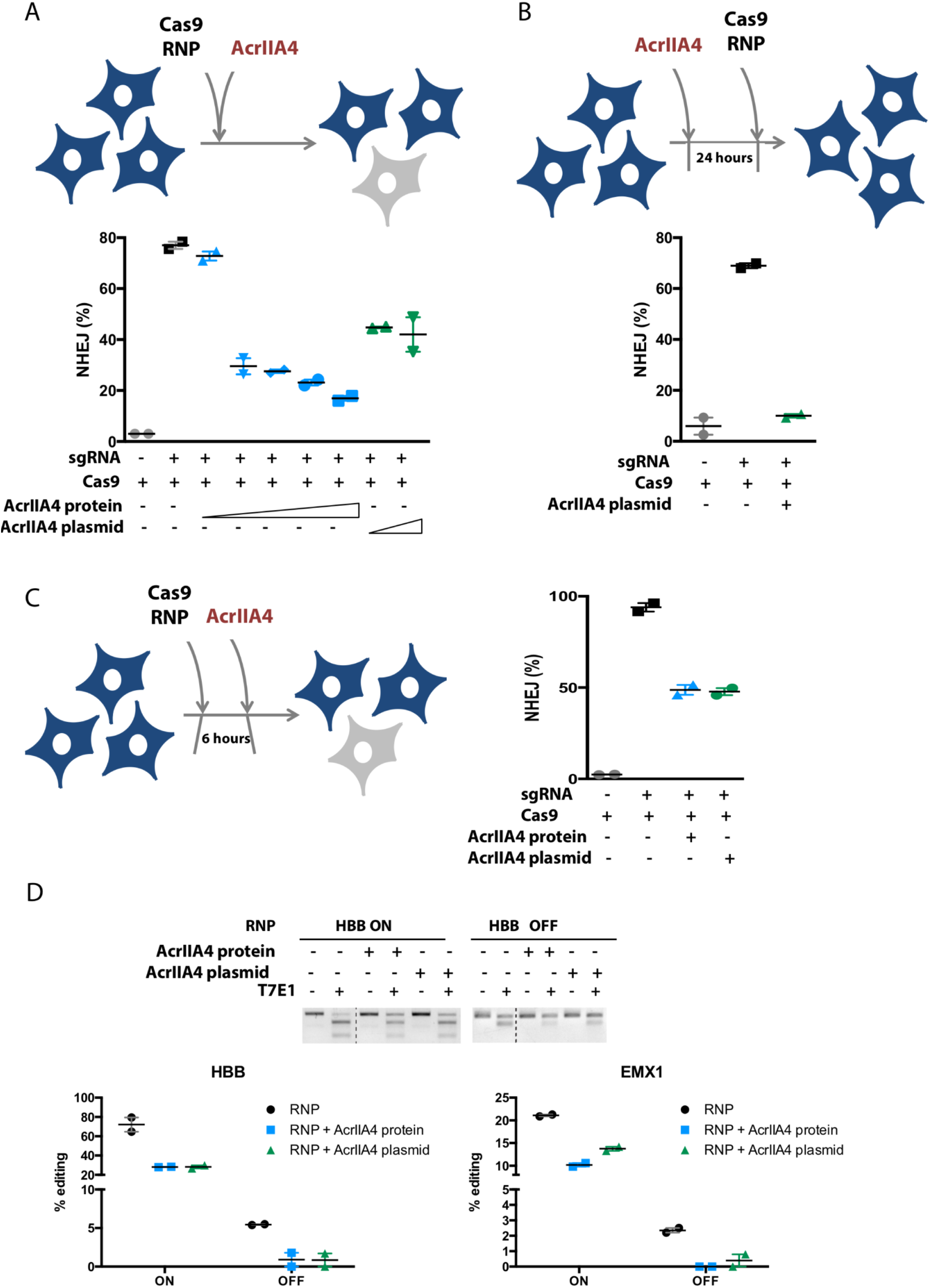
Timed delivery of AcrIIA4 differentially inhibits on- and off-target genome editing in human cells. A) Simultaneous delivery of Cas9 RNP and AcrIIA4 inhibits Cas9-mediated gene targeting in human cells. K562 cells with a chromosomally integrated BFP (BFP-K562) were nucleofected with Cas9 RNP and AcrIIA4 protein (Cas9:AcrIIA4 (molar ratio), 1:0.5, 1:1, 1:2, 1:3, 1:5) or plasmid (0.7μg and 2.8μg). NHEJ frequencies are quantified by loss of BFP expression in BFP-K562 cells 96 hours post nucleofection via flow cytometry. Data presented as mean ± SEM from at least two biological replicates. B) Administration of AcrIIA4 prior to Cas9 RNP completely inhibits Cas9-mediated gene targeting. BFP-K562 cells were nucleofected with AcrIIA4 plasmid (0.7μg) 24 hours prior to Cas9 RNP delivery. Data presented as mean ± SEM from at least two biological replicates. C) Delivery of AcrIIA4 after introduction of Cas9 RNP yields intermediate inhibition of Cas9 activity. BFP-K562 cells were nucleofected with AcrIIA4 protein (Cas9:AcrIIA4 (molar ratio), 1:5) or plasmid (0.7μg) 6 hours post Cas9 RNP delivery. Data presented as mean ± SEM from at least two biological replicates. D) Proper timing of AcrIIA4 delivery diminishes off-target editing events while largely retaining on-target editing. K562 cells were nucleofected with either HBB or EMX1 targeting Cas9 RNP 6 hours prior to AcrIIA4 protein (Cas9:AcrIIA4 (molar ratio), 1:5) or plasmid (0.7μg) delivery. (Top) Representative T7 endonuclease I assay for visualization of HBB on- and off-target editing. (Bottom) Quantification of on- and off-target editing at HBB and EMX1, as measured by TIDE analysis.

Using a given guide RNA, Cas9 may target both an on-target site and off-target loci (9-12). But several lines of evidence suggest that off-target sites may be bound without being immediately cleaved (13-15). Cas9 displaced from uncleaved sites (e.g. by cellular factors) would be available for inhibition by AcrIIA4. Since inhibitor timing experiments suggested that at least 50% of on-target Cas9 gene editing takes place within the first 6 hours (Fig. 4C), we asked whether off-target editing could be reduced by properly timed addition of inhibitor.

We examined on- and off-target editing using guide RNAs targeting the EMX1 and HBB loci. The off-target sites for both the EMX1 and HBB guides have been previously described (10, 16, 17). Notably, the HBB guide is of therapeutic interest to edit the causative mutation of sickle cell disease, but has an off-target that is nearly identical to the on-target site. We edited the EMX1 and HBB loci using Cas9 RNPs with and without pre-introduction of AcrIIA4 protein or plasmid. Consistent with experiments at the BFP locus, we found that timed addition of AcrIIA4 retained substantial levels of on-target editing. Strikingly, both T7E1 and TIDE analysis showed that timed addition of AcrIIA4 almost completely abolished off-target editing at both HBB and EMX1 (Fig. 4D). This suggests that Cas9 inhibitors have the potential to be useful in reducing off-target events during research and therapeutic applications.

The recent and rapid expansion of the Cas9 toolkit for gene editing applications has lacked an inducible “off-switch” to prevent undesired gene editing. Newly discovered protein inhibitors, encoded by bacteriophages, provide an attractive solution to this problem as these proteins are small and function well in human cells (3). Here, we demonstrate that AcrIIA4, the most potent SpyCas9 inhibitor in human cells, acts as a DNA mimic to block PAM recognition. The deployment of DNA mimics to inhibit a DNA-binding protein is an elegant solution that is often deployed in the phage-host arms race. A recent structural study revealed that a Class 1 anti-CRISPR (AcrF2) uses a similar strategy (18). Further, phage-encoded restriction enzyme inhibitor proteins have long been known to mimic the DNA target (19). Our results further show that Cas9 inhibitors delivered as either protein or expressed from a plasmid can modulate the efficacy of gene editing at multiple loci in human cells. Although pre-addition of inhibitor almost completely abolishes overall gene editing, timed addition of inhibitor after initiating Cas9-sgRNA-based gene editing can adjust the amount of time that Cas9 is active in the nucleus, thereby selectively limiting off-target editing. We anticipate that Cas9 inhibitors could be broadly useful in situations where precise control of either on- or off-target gene editing is desirable, such as during allele-specific therapeutic editing.

## Acknowledgments

EM-derived maps and atomic coordinates of the Cas9-sgRNA-AcrIIA4 structure have been deposited in the EM Databank (EMDB) and the Protein Data Bank (PDB) with accession codes **EMD-XXXX and YYYY, respectively.** The EM data was collected in the EM facility of HHMI Janelia Research Campus. We thank Rick Huang and Zhiheng Yu for expert electron microscopy assistance; and members of the Bondy-Denomy, Corn, Doudna and Nogales labs for helpful discussions and critical reading of the manuscript. F.J. is a Merck Fellow of the Damon Runyon Cancer Research. J.E.C. is supported by the Li Ka Shing Foundation and the Heritage Medical Research Institute. J.S. is supported by the National Institute on Aging of the National Institutes of Health under Award Number T32 AG000266. J.B.-D. and B.R. are supported by the University of California San Francisco Program for Breakthrough in Biomedical Research, funded in part by the Sandler Foundation, and an NIH Office of the Director Early Independence Award (DP5-OD021344). J.A.D. and E.N. are Investigators of the Howard Hughes Medical Institute. This work was supported in part by HHMI. The content is solely the responsibility of the authors and does not necessarily represent the official views of the National Institutes of Health.

